## Supplementary for "An Immunological Synapse Formation Between T Regulatory Cells and Cancer-Associated Fibroblasts Promotes Tumor Development"

**Supplementary Information**

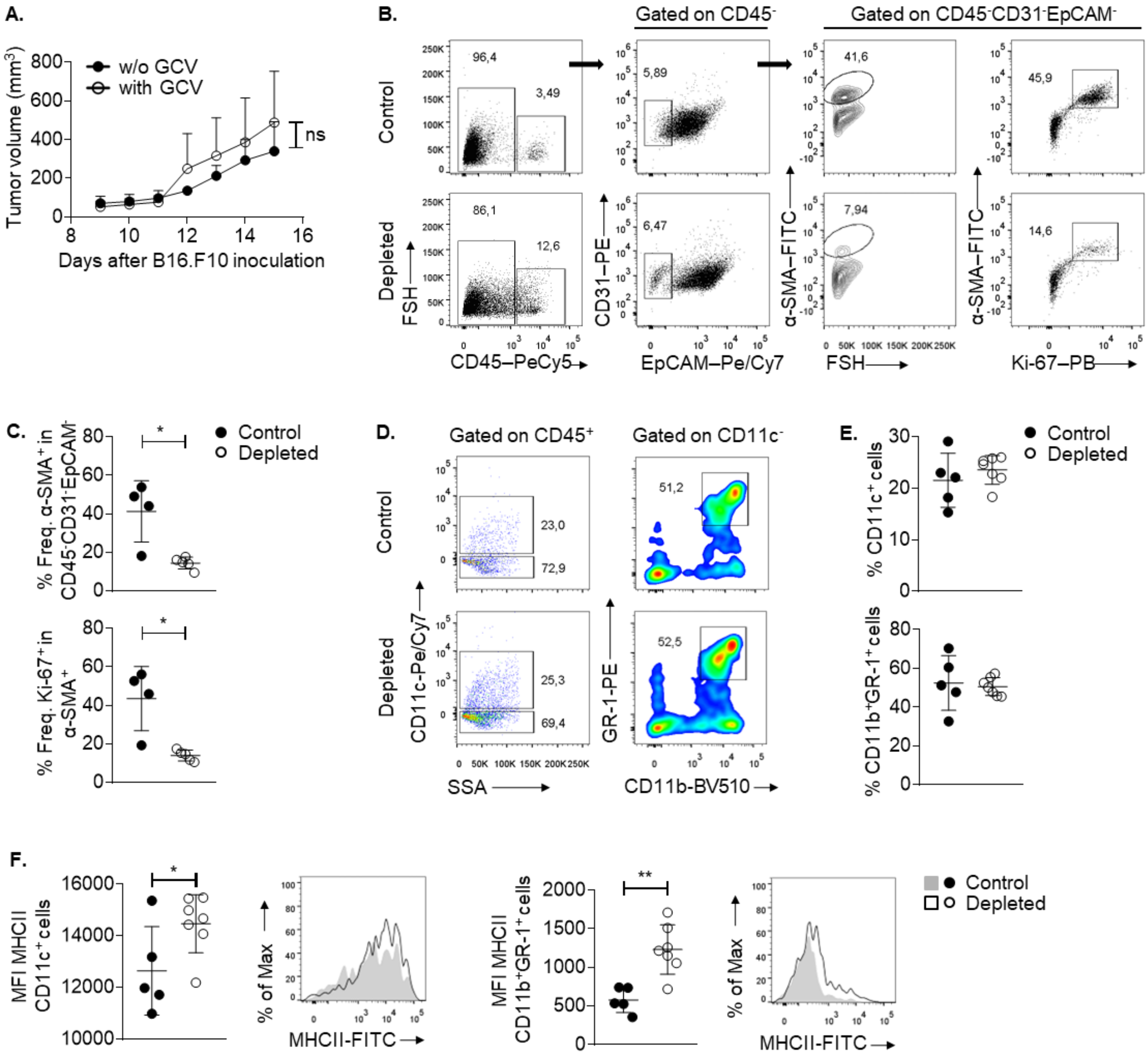

**Supplementary Figure 1. Ablation of αSMA$^+$ CAFs increases the immunogenicity of tumor infiltrating myeloid cells.** **(A)** Tumor volume (mm$^3$) of B16.F10 inoculated PBS-treated (w/o GCV, n=3) and GCV-treated (with GCV, n=4) C57BL/6 mice. **(B)** Representative FACS plots and percentages of intra-tumoral αSMA$^+$ cells on Day15 after B16.F10 inoculation of PBS-treated (control, n=3) and GCV-treated (depleted, n=4) αSMA-tk$^+$ mice. **(C)** Representative FACS plots and percentages of intra-tumoral CD11c$^+$ and CD11b$^+$GR-1$^+$ cells on Day15 after B16.F10 inoculation of PBS-treated (control, n=5) and GCV-treated (depleted, n=7) αSMA-tk$^+$ mice **(D)**

Representative overlays and MHC II mean fluorescence intensity (MFI) of intra-tumoral $CD11c^+$ and $CD11b^+GR\text{-}1^+$ cells on Day15 after B16.F10 inoculation of PBS-treated (control, n=5) and GCV-treated (depleted, n=7) $\alpha SMA\text{-}tk^+$ mice. Data are shown as mean ± SD. Representative data from four (a, b) and two (C, D) independent experiments are shown. Unpaired two-tailed t-test (A-D). *$P<0.05$, **$P<0.01$, ns=non-significant. n=biologically independent samples.

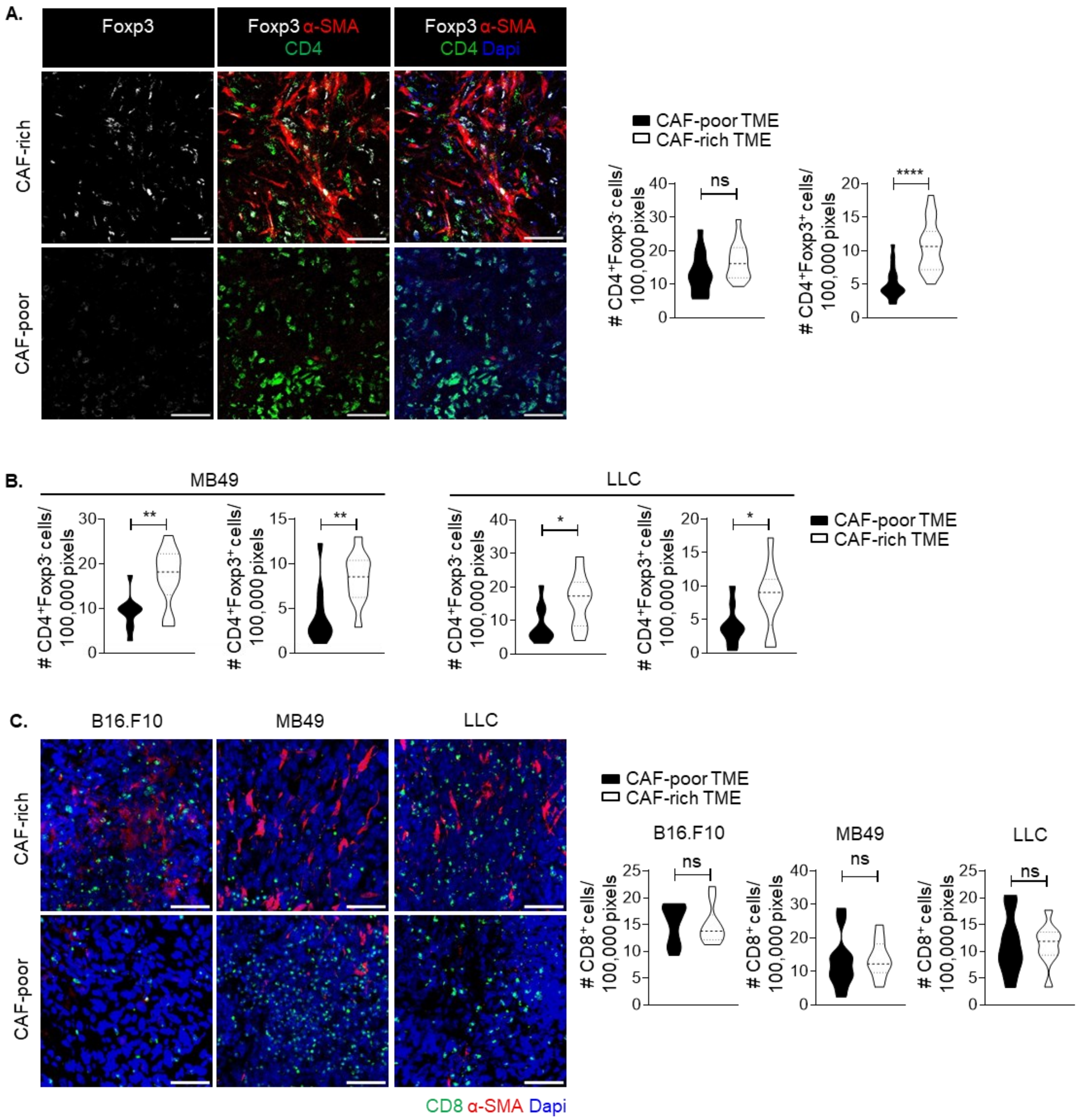


**Supplementary Figure 2. T effector cells (CD4$^+$Foxp3$^-$) and Treg cells (CD4$^+$Foxp3$^+$) accumulate in α-SMA$^+$ CAF-rich areas while CD8$^+$ T cells are equally distributed in α-SMA$^+$ CAF-rich and -poor areas of the TME**. **(A)** Representative images of immunofluorescence confocal microscopy for α-SMA-RFP (red), Foxp3 (silver white), and CD4 (green) in B16.F10 melanoma cryosections isolated from αSMARFP mice. Scale bar: 50μm. Quantitation plots depicting numbers of CD4$^+$Foxp3$^-$ cells and CD4$^+$Foxp3$^+$ cells in CAF-rich and CAF-poor regions.

8 fields or more per tumor were captured. **(B)** Quantitation plots depicting numbers of $CD4^{+}Foxp3^{-}$ cells and $CD4^{+}Foxp3^{+}$ cells in CAF-rich and CAF-poor regions. 8 fields or more per tumor were captured. **(C)** Representative images of immunofluorescence confocal microscopy for α-SMA-RFP (red) and CD8 (green) in B16.F10, MB49 and Lewis Lung Carcinoma tumor cryosections isolated from $\alpha SMA^{RFP}$ mice. Scale bar: 100µm. Quantitation plots depicting numbers of $CD8^{+}$ cells in CAF-rich and CAF-poor regions. 8 fields or more per tumor were captured. Paired two-tailed t-test (A-C); *P<0.05, **P<0.01, ****P<0.001, ns=not significant.

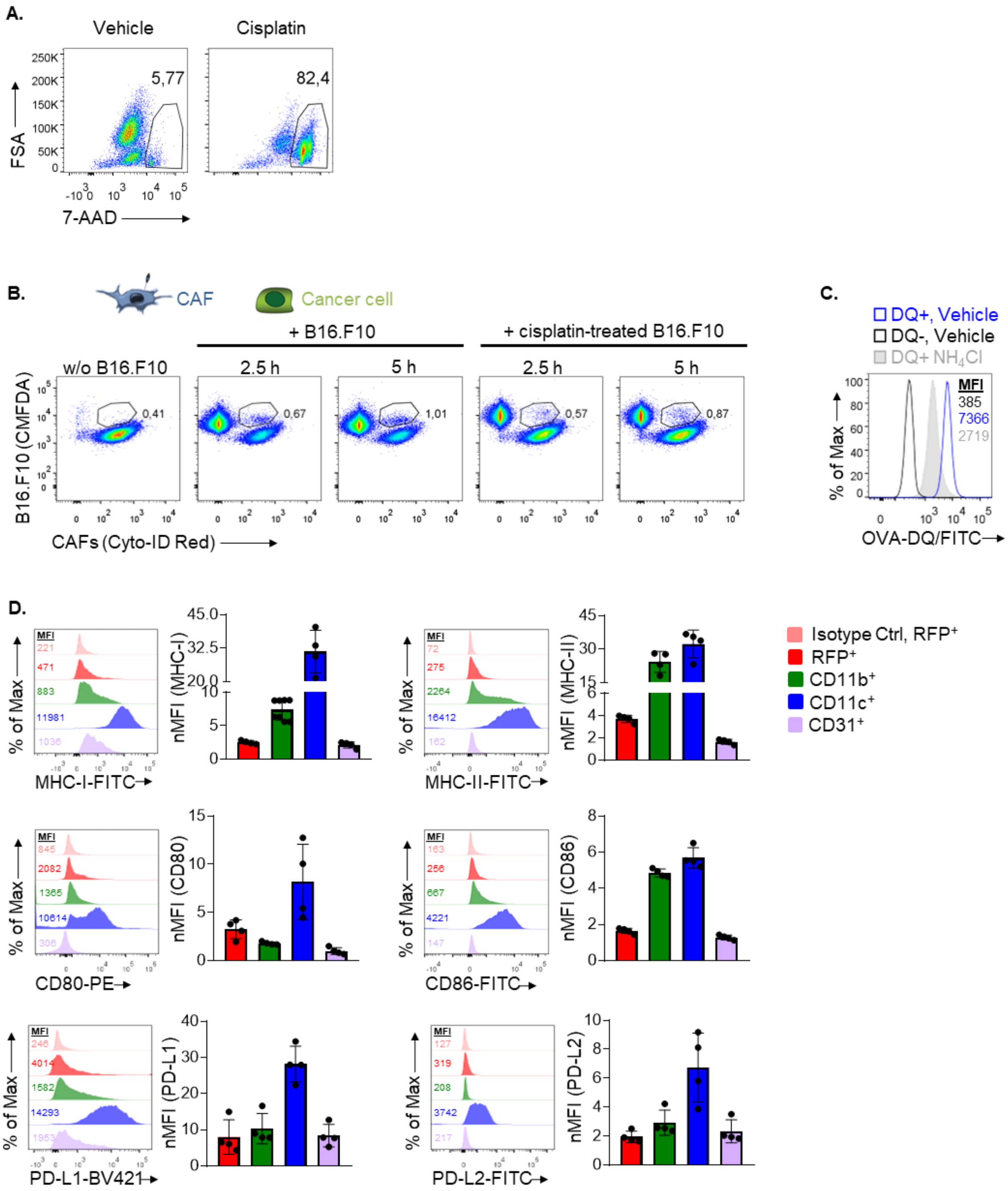


**Supplementary Figure 3. α-SMA$^{+}$ CAFs have the capacity to uptake, process and present tumor antigens.** (A) Representative FACS plots showing 7-Aminoactinomycin D (7-aad) staining in cisplatin-treated or vehicle-treated B16.F10 cells. **(B)** CYTO-ID Red cell tracer-labelled CAFs were co-cultured with CMFDA-stained B16.F10 or cisplatin-treated apoptotic B16.F10 cells for 2.5h and 5h. Representative flow cytometry plots depicting percentages of CAFs harboring cancer cells. **(C)** Representative overlay, mean fluorescence intensity (MFI) depicting processing of DQ-OVA by $NH_4Cl$-treated CAFs and vehicle-treated CAFs. **(D)** Representative overlays and normalized MFIs (nMFIs) of MHC-I, MHC-II, CD80, CD86, PD-L1, PD-L2 in intra-tumoral $CD45^+CD11b^+$ cells, $CD45^+CD11c^+$ cells, $CD45^-CD31^+$ cells and $CD45^-CD31^-$α-$SMA^+$ CAFs on Day15 after B16.F10 inoculation of $αSMA^{RFP}$ mice (n=4). Data are shown as mean ± SD. Representative data from three (B, C), two (D) independent experiments are shown.

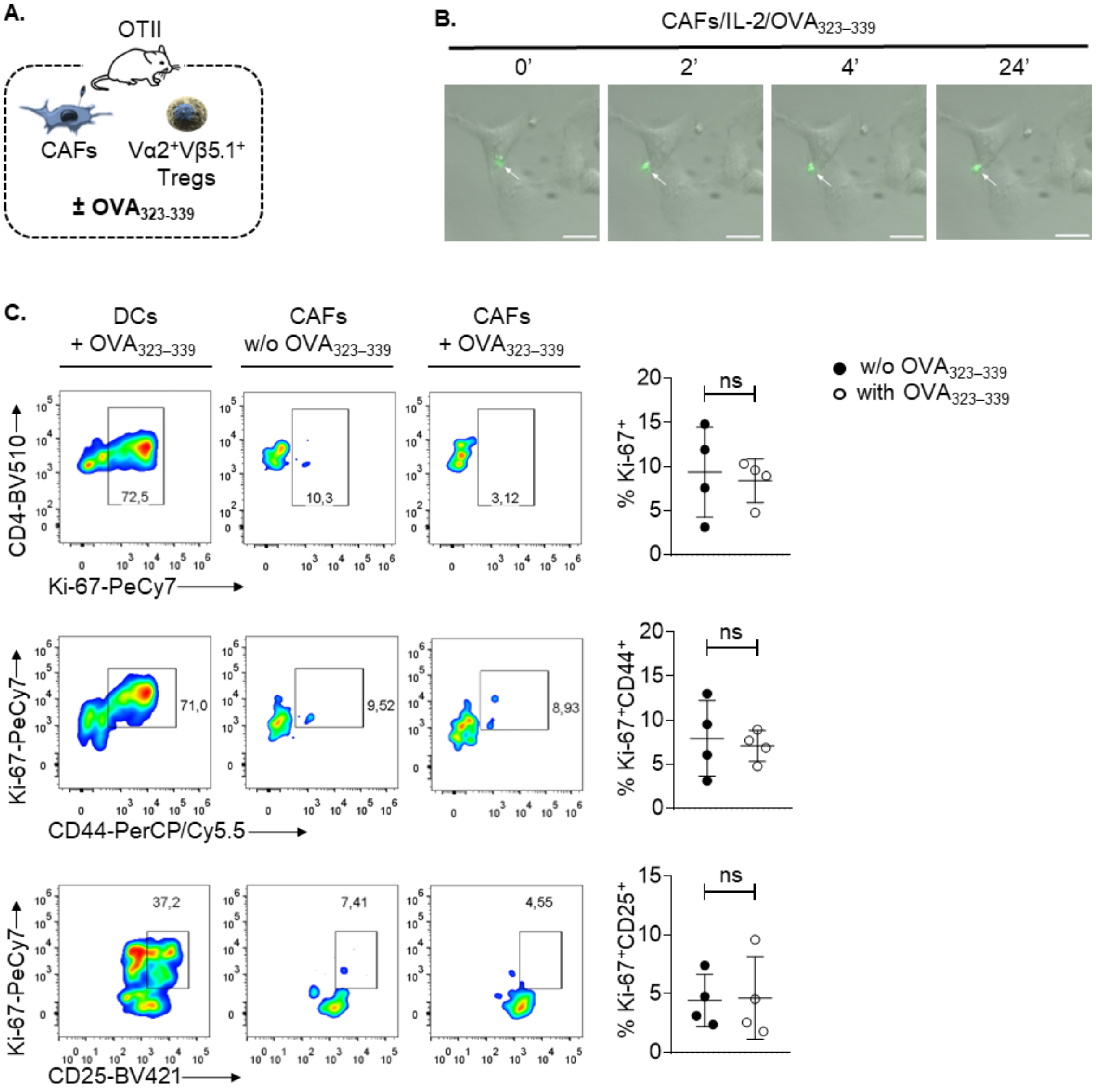


**Supplementary Figure 4. α-SMA⁺ CAFs form prolonged interactions with Treg cells in an antigen-specific fashion, yet fail to activate T effector cells. (A)** Schematic representation depicting the co-culture setup used in OTII experiments. **(B)** CMFDA-stained Treg cells isolated from total lymph nodes and spleen of OTII mice were co-cultured with CAFs in the presence of IL-2 and $OVA_{323\text{–}339}$. CAF-Treg cell interactions were monitored with time-lapse microscopy (4h, 5'/frame). Scale bar: 25μm. **(C)** Flow cytometry plots (left), percentages (right) of Ki-67$^+$,

$CD25^+Ki\text{-}67^+$ and $CD25^+Ki\text{-}67^+$ OTII T effector cells following co-culture with DCs or CAFs in the absence or presence of $OVA_{323\text{-}339}$ peptide. Data are shown as mean ± SD. Representative data from two (B, C) independent experiments are shown. Unpaired two-tailed t-test (C). ns=non-significant.

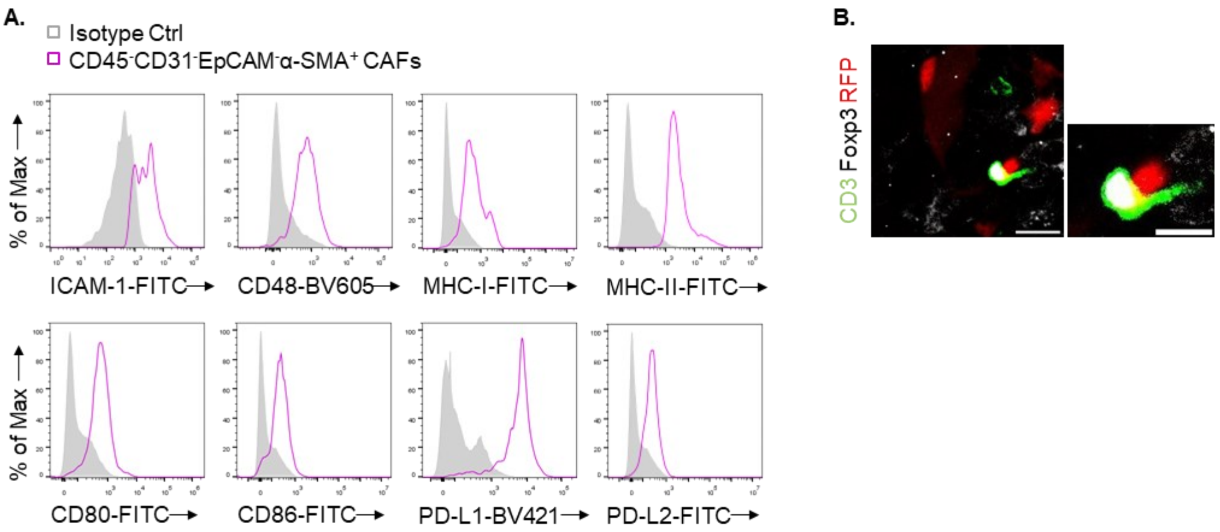


**Supplementary Figure 5. A-SMA$^{+}$ CAFs can form synapses with Treg cells in an antigen-specific manner. (A)** Representative overlays of CD45$^{-}$CD31$^{-}$EpCAM$^{-}$α-SMA$^{+}$ CAFs isolated from Day15 B16.F10 tumors, stained for ICAM-1, CD48, MHC-I, MHC-II, CD80, CD86, PD-L1, PD-L2 vs isotype control Ab. **(B)** Representative images of αSMA-RFP (red), Foxp3 (silver white), CD3 (green) immunofluorescence from Day15 B16.F10 (n=4) tumor cryosections derived from αSMA$^{RFP}$ mice. Scale bars: 15μm, 9μm (inset). Representative data from two (A), three (B) independent experiments are shown. n=biologically independent samples.

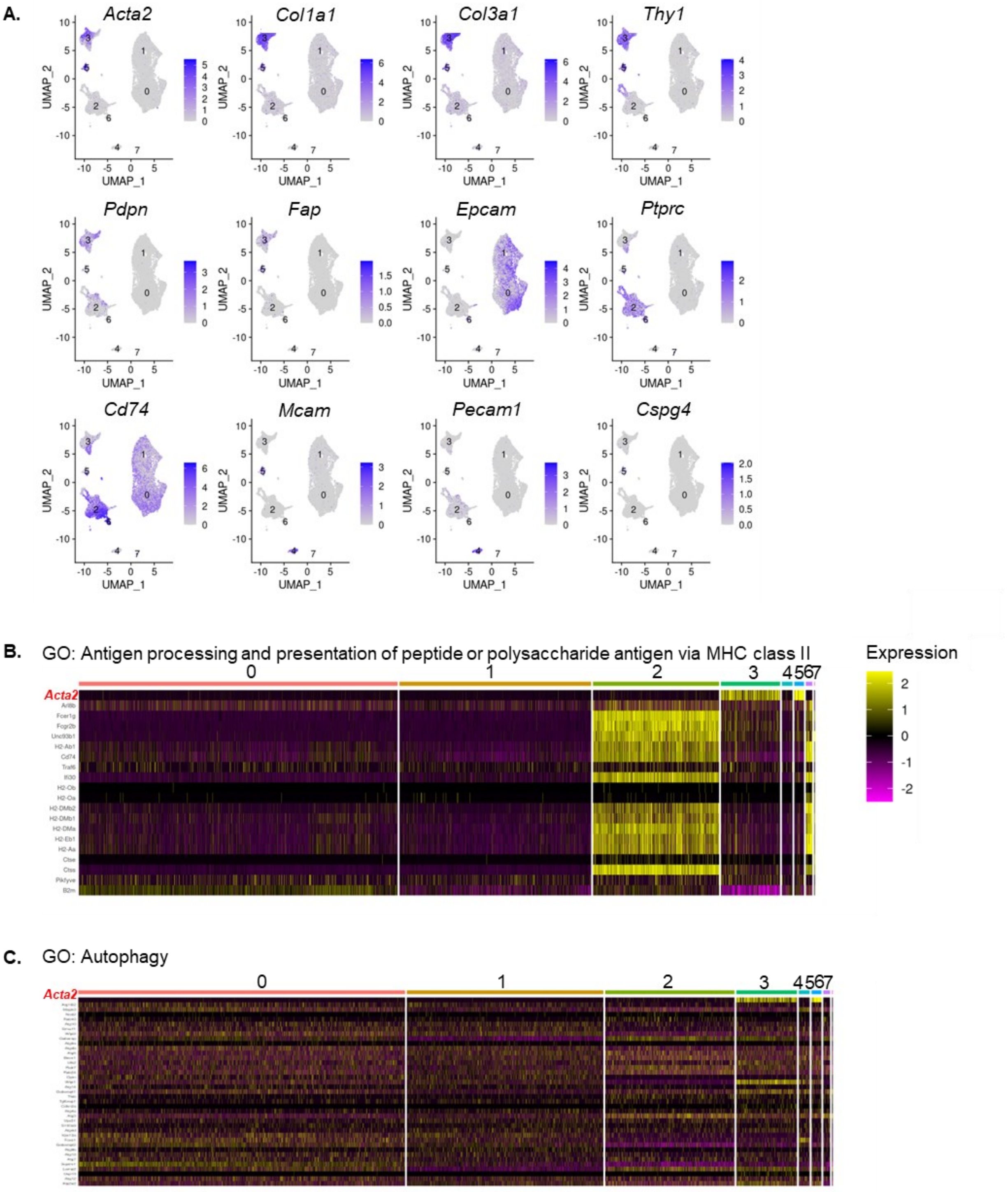


**Supplementary Figure 6. α-SMA⁺ CAFs upregulate autophagy in 4T1 breast tumors. (A)** UMAP plots of *Acta2, Col1a1, Col3a1, Thy1, Pdpn, Fap, Epcam, Ptprc, Cd74, Mcam, Pecam1, Cspg4* genes from Grauel *et al.* data. Color keys indicate log-normalized expression. **(B)** Heatmap displaying genes associated with GO: 0002504, antigen processing and presentation of peptide or polysaccharide antigen via MHC class II for all identified clusters from Grauel *et al.* data. **(C)** Heatmap displaying genes associated with GO: 0006914, autophagy for all identified clusters from Grauel *et al.* data.

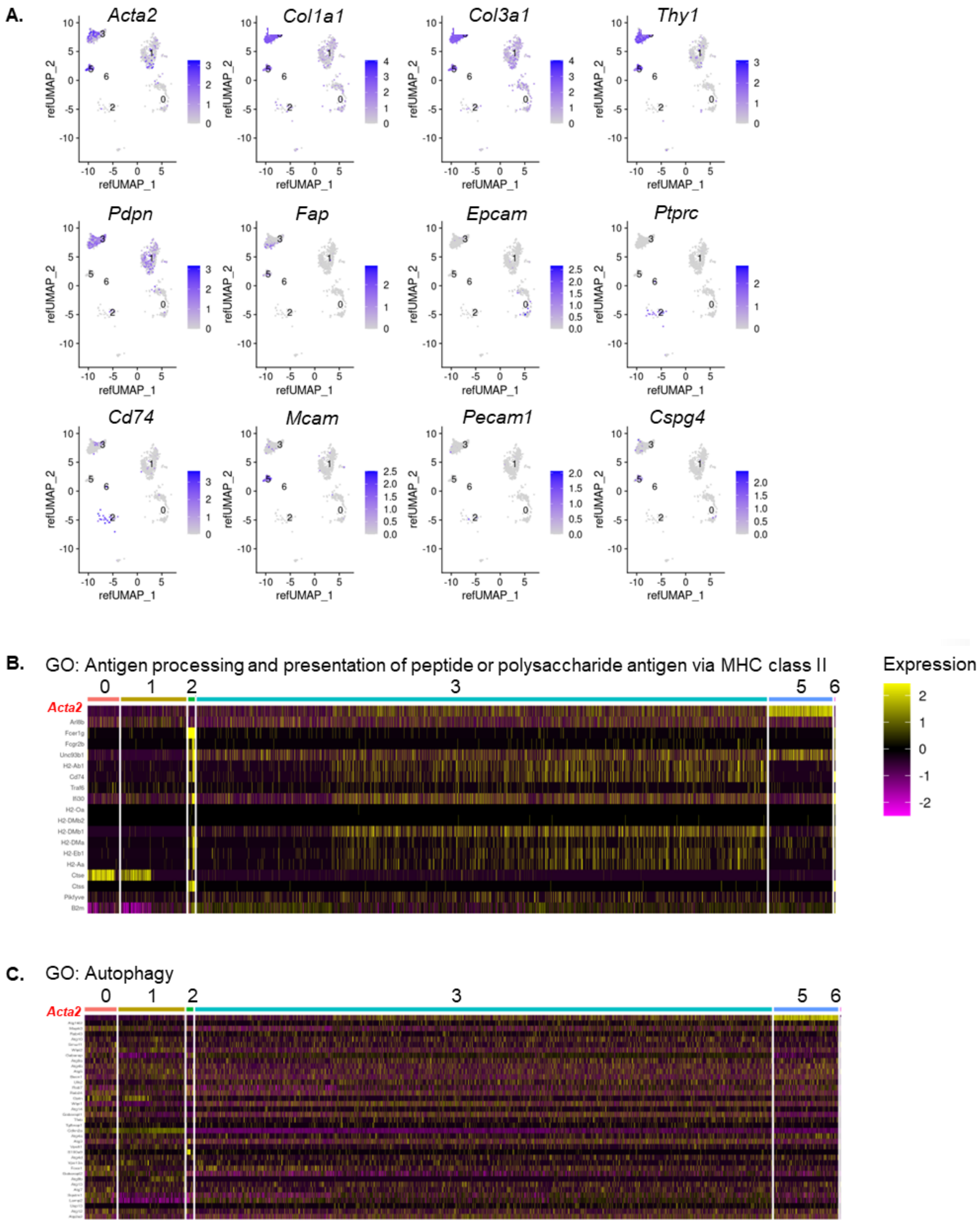


**Supplementary Figure 7. α-SMA⁺ CAFs upregulate autophagy in KPC pancreatic tumors. (A)** UMAP plots of *Acta2, Col1a1, Col3a1, Thy1, Pdpn, Fap, Epcam, Ptprc, Cd74, Mcam, Pecam1, Cspg4* genes from Elyada *et al.* data. Color keys indicate log-normalized expression. **(B)**

Heatmap displaying genes associated with GO: 0002504, antigen processing and presentation of peptide or polysaccharide antigen via MHC class II for all identified clusters from Elyada *et al.* data. **(C)** Heatmap displaying genes associated with GO: 0006914, autophagy for all identified clusters from Elyada *et al.* data.

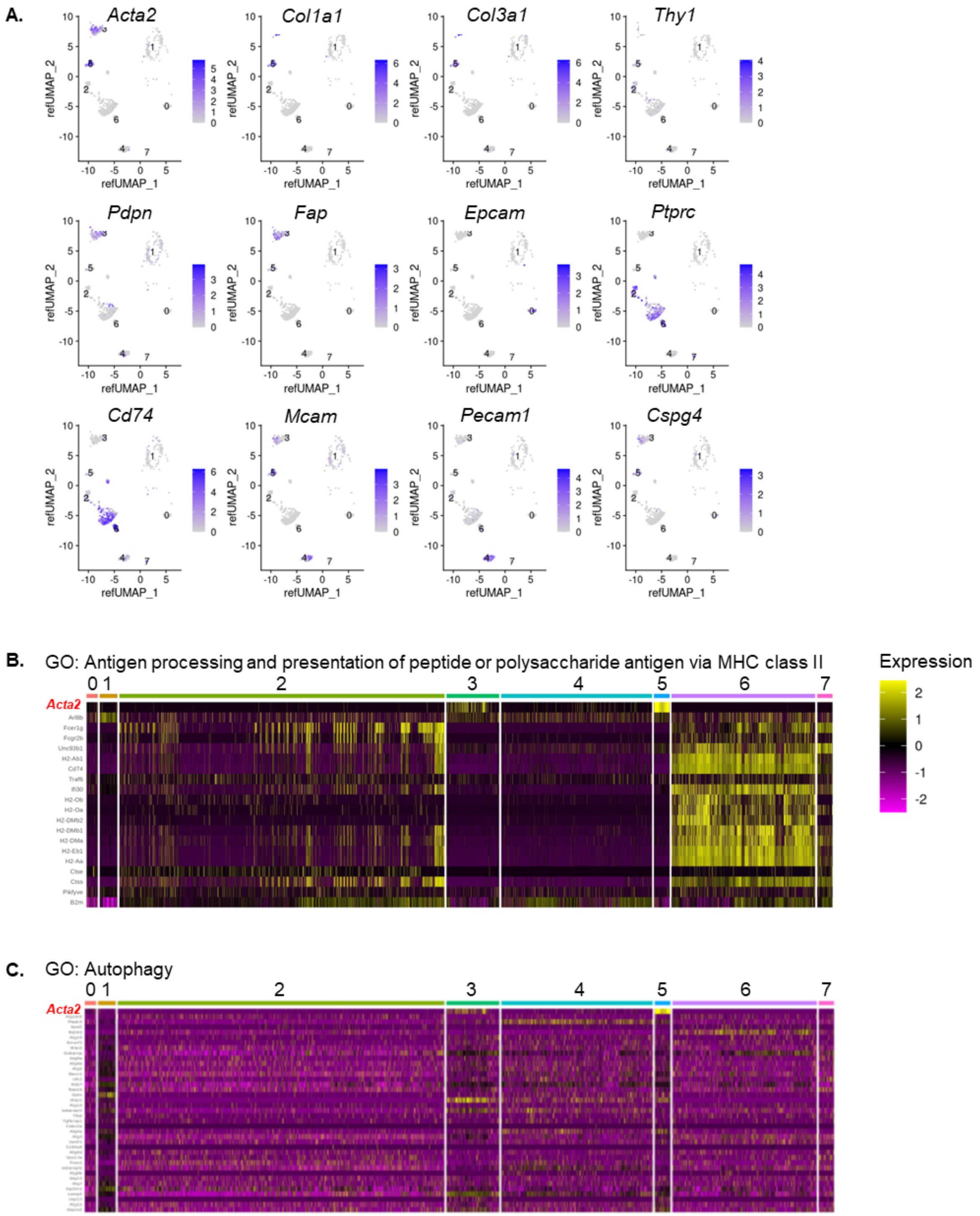


**Supplementary Figure 8. α-SMA$^{+}$ CAFs upregulate autophagy in B16.F10 melanomas. (A)** UMAP plots of *Acta2, Col1a1, Col3a1, Thy1, Pdpn, Fap, Epcam, Ptprc, Cd74, Mcam, Pecam1, Cspg4* genes from Davidson *et al.* data. Color keys indicate log-normalized expression. **(B)**

Heatmap displaying genes associated with GO: 0002504, antigen processing and presentation of peptide or polysaccharide antigen via MHC class II for all identified clusters from Davidson *et al.* data. **(C)** Heatmap displaying genes associated with GO: 0006914, autophagy for all identified clusters from Davidson *et al.* data.

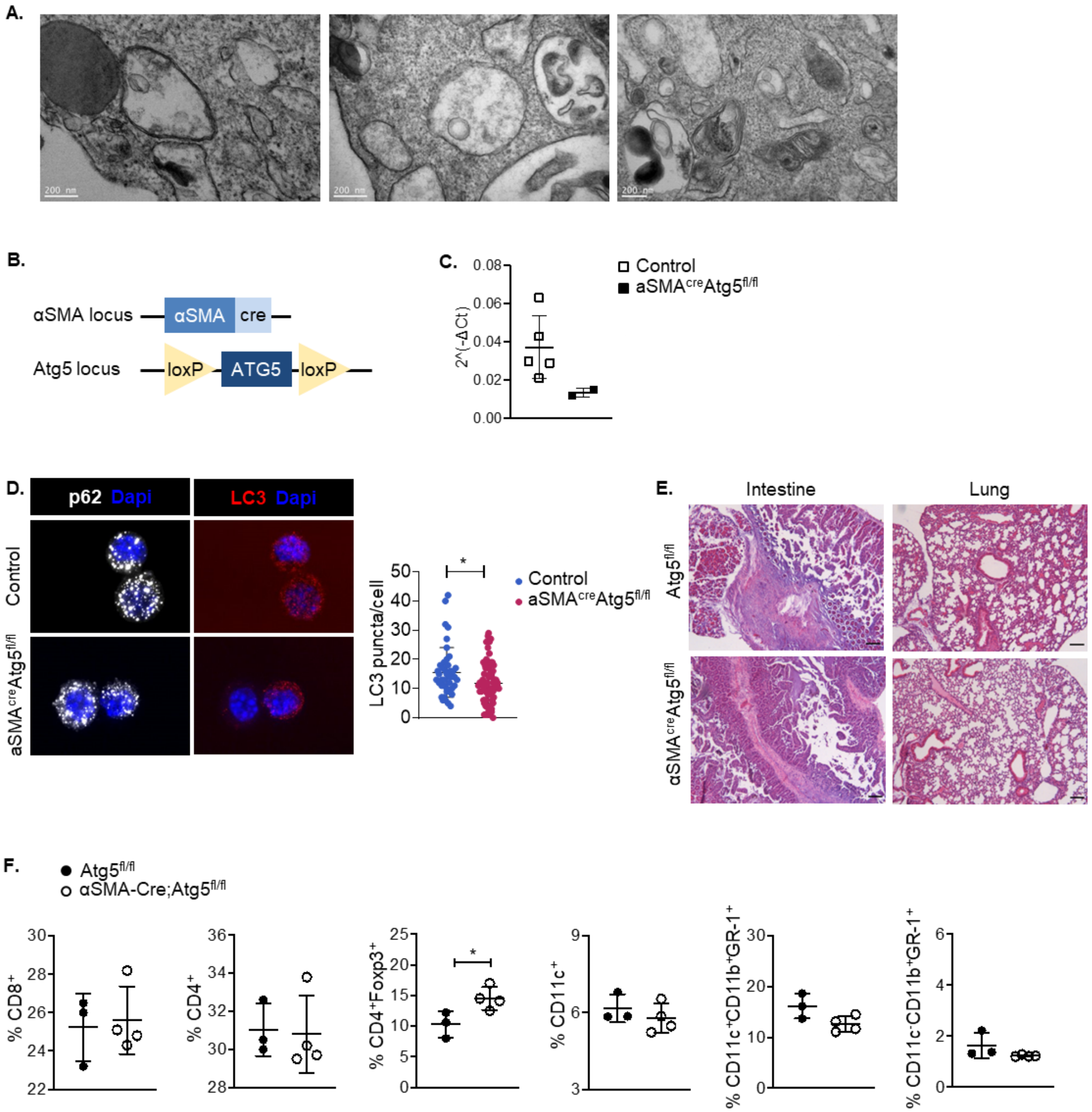


**Supplementary Figure 9. Autophagy deficiency in αSMA$^{+}$ CAFs does not affect the development of immune cells in a naive state. (A)** Transmission electron microscopy images of isolated CAFs. Scale bar: 200nm as indicated in figure. **(B)** αSMA$^{cre}$Atg5$^{fl/fl}$ mice, schematic illustration. Mice were generated with the cre-LoxP system, by crossing αSMA$^{cre}$ mice with Atg5$^{fl/fl}$ mice where cre recombinase is expressed under the control of the *Acta2* gene promoter and the exon 3.1 of ATG5 gene is flanked by the LoxP sites containing neomycin resistant cassette.

αSMA$^{cre}$Atg5$^{fl/fl}$ mice have impaired autophagy in αSMA-expressing cells. **(C)** Expression ($2^{-\Delta Ct}$) of *Atg5* mRNA in isolated CAFs from control (n=5) and aSMA$^{cre}$Atg5$^{fl/fl}$ (n=2) mice, analyzed by quantitative real time RT-PCR. **(D)** Representative images of immunofluorescence confocal microscopy for LC3 (red) and p62 (silver white) in CAFs isolated from control (n=42) and aSMA$^{cre}$Atg5$^{fl/fl}$ (n=72) mice. Scale bar: 12μm. LC3 puncta/cell are depicted. **(E)** Hematoxylin-and-eosin staining of intestine and lungs from steady state Atg5$^{fl/fl}$ and αSMA$^{cre}$Atg5$^{fl/fl}$ mice aged 8mo. Original magnification ×20. Scale bar: 10μm. **(F)** Percentages of CD4$^{+}$ T cells, CD8$^{+}$ T cells and Treg cells (CD4$^{+}$Foxp3$^{+}$) in total lymph nodes and DCs (CD11c$^{+}$), inflammatory DCs (CD11c$^{+}$CD11b$^{+}$GR-1$^{+}$) and MDSCs (CD11c$^{-}$CD11b$^{+}$GR-1$^{+}$) in spleens from steady state Atg5$^{fl/fl}$ and αSMA$^{cre}$Atg5$^{fl/fl}$ aged 8mo. Representative data from two (E, F) independent experiments are shown. Unpaired two-tailed t-test (F), *P<0.05, n=biologically independent samples.

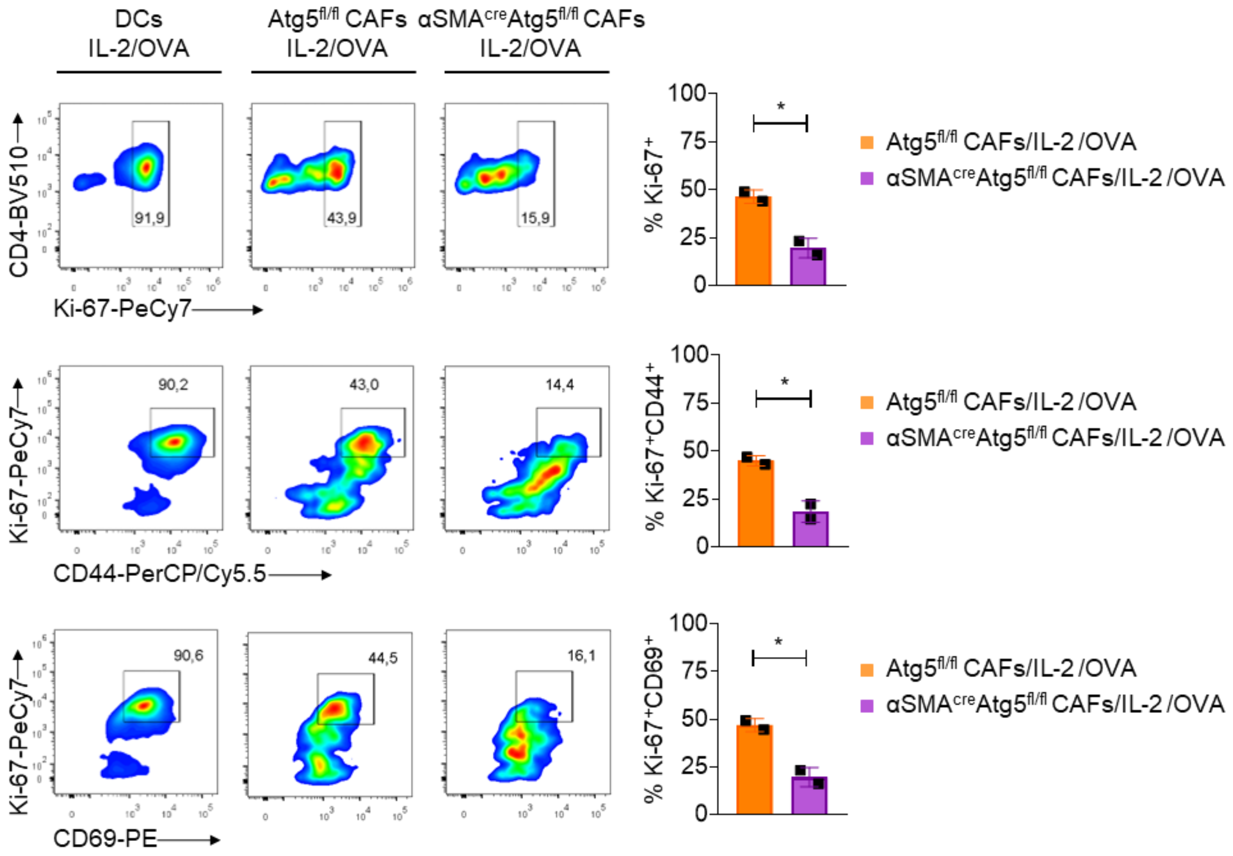


**Supplementary Figure 10. α-SMA$^+$ CAFs with depleted autophagy display diminished capacity to process the entire antigen and to activate T regulatory cells**. Flow cytometry plots (left), percentages (right) depicting expression of CD4, Ki-67, CD44 and CD69 in OTII CD4$^+$Foxp3$^+$ Treg cells co-cultured with isolated DCs or CAFs from Atg5$^{fl/fl}$ mice (n=2) or CAFs from αSMA$^{cre}$Atg5$^{fl/fl}$ mice (n=2), in the presence of IL-2 and whole OVA protein. Data are shown as mean ± SD. Representative data from two independent experiments are shown. *P<0.05, n=biologically independent samples.

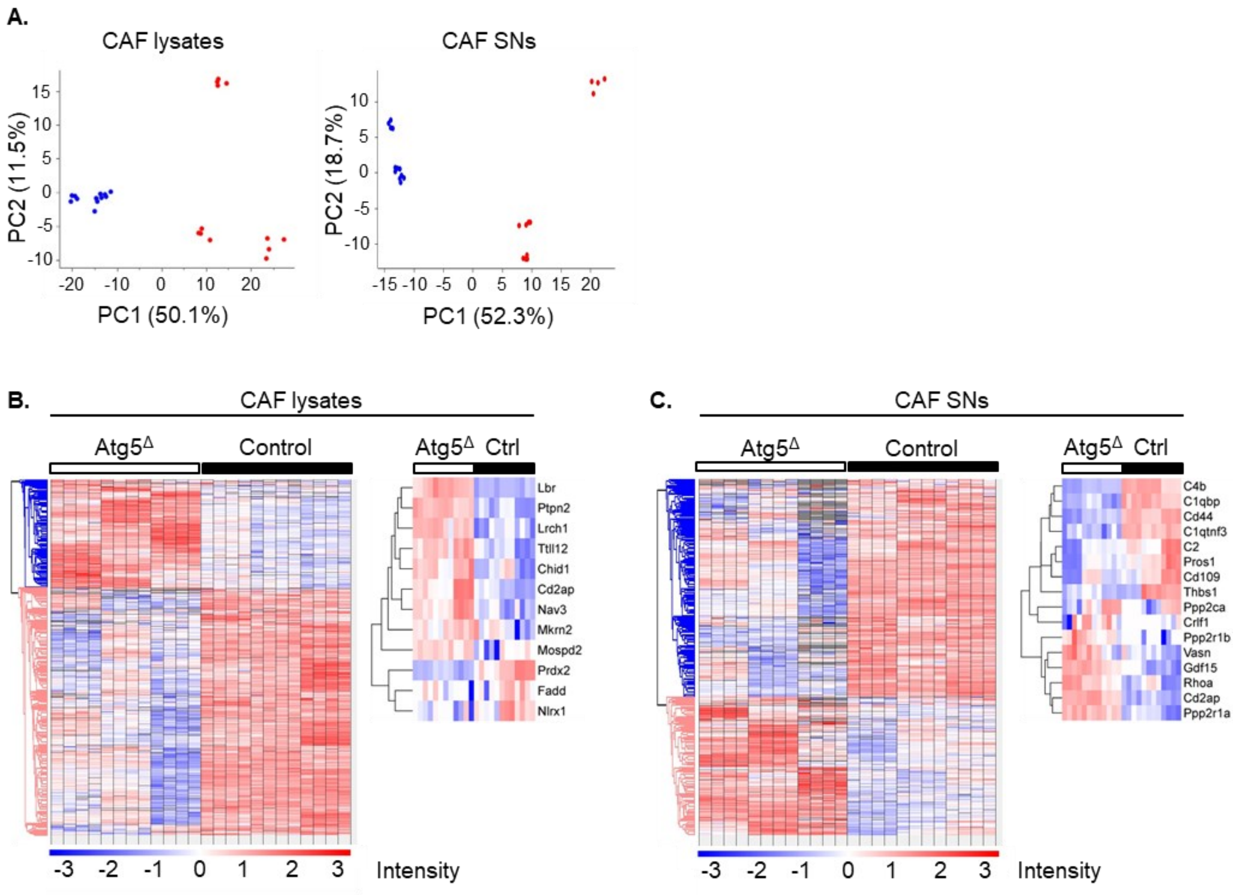


**Supplementary Figure 11. Autophagy deficient CAFs are more abundant in proteins associated with inflammatory processes. (A)** PCA plots of samples used for proteomic analysis of CAF lysates (left) and supernatants (SNs, right). **(B-C)** Heatmaps with scaled expression values (row z-score) of all differentially expressed proteins (DEPs, left) and DEPs associated with inflammatory responses (right) between CAFs lysates **(B)** or CAF supernatants **(C)** isolated from control (n=3) and aSMA$^{cre}$Atg5$^{fl/fl}$ (n=3) mice.

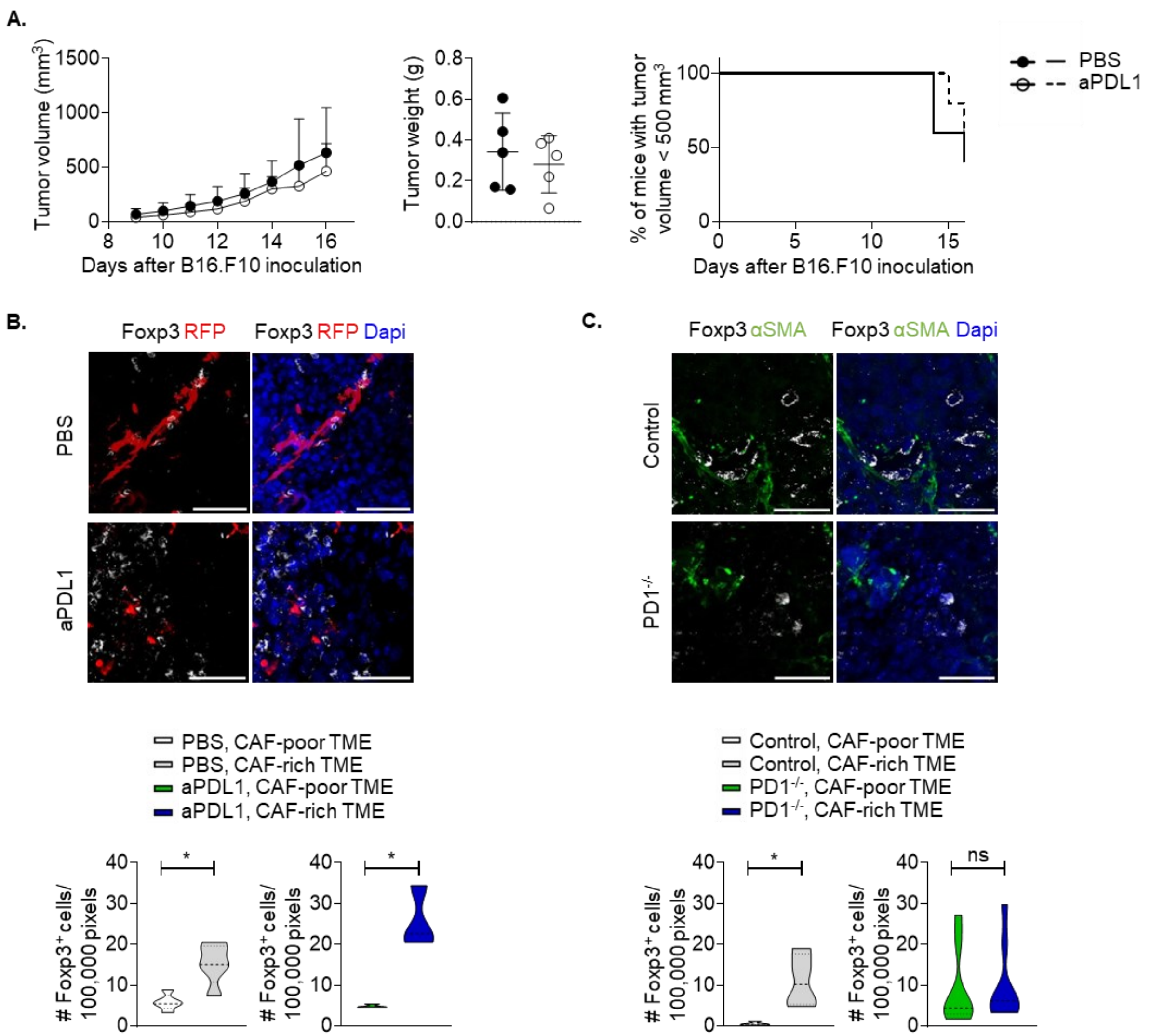


**Supplementary Figure 12. PD-L1 blockade does not disrupt the synaptic formation between αSMA$^+$ CAFs and Foxp3$^+$ Treg cells in the TME.** (A) Tumor volume (mm$^3$), tumor weight (g) and percentage of mice bearing tumors <500 mm$^3$ of B16.F10 inoculated PBS-treated (PBS, n=5) and anti-PD-L1-treated (aPD-L1, n=5) αSMA$^{RFP}$ mice. (B) Representative images of αSMA-RFP, Foxp3 immunofluorescence from Day15 B16.F10 tumor cryosections derived from PBS-treated (PBS, n=3) and anti-PD-L1-treated (aPD-L1, n=3) αSMA$^{RFP}$ mice. 5 fields or more per tumor were captured. Scale bar: 40μm. Quantitation plots depicting number of Foxp3$^+$ cells infiltrating Day15 B16.F10 tumors in CAF-poor and CAF-rich regions of PBS-treated (PBS, n=3) and anti-PD-L1-

treated (aPD-L1, n=3) αSMA$^{RFP}$ mice. (C) Representative images of αSMA, Foxp3 immunofluorescence from Day15 B16.F10 tumor cryosections derived from control (n=5) and PD-1$^{-/-}$ (n=3) mice. 5 fields or more per tumor were captured. Scale bar: 40μm. Quantitation plots depicting number of Foxp3$^{+}$ cells infiltrating Day15 B16.F10 tumors in CAF-poor and CAF-rich regions of control (n=5) and PD-1$^{-/-}$ (n=3) mice. Data are shown as mean ± SD. Representative data from four (A), three (B), two (C) independent experiments are shown. Unpaired two-tailed t-test (A), two-tailed log(rank) test (A), Paired t-test (B, C). *P<0.05, **P<0.01, ns=non-significant. n=biologically independent samples.

**Movie S1.** Representative time-lapse video microscopy showing interaction between CAFs and $CD4^{+}CD25^{+}GITR^{+}V\alpha2^{+}V\beta5.1^{+}$ Tregs sorted from OTII mice, in the presence of $OVA_{323\text{-}339}$. Related to Figure 3. 4fps, scale bar: 100 pixels.

**Movie S2.** Representative time-lapse video microscopy showing interaction between CAFs and $CD4^{+}CD25^{+}GITR^{+}V\alpha2^{+}V\beta5.1^{+}$ Tregs sorted from OTII mice, in the absence of $OVA_{323\text{-}339}$ (vehicle). Related to Figure 3. 4fps, scale bar: 100 pixels.

**Movie S3.** Representative video from 3D fluorescent confocal microscopic imaging, showing interaction between CAFs and $CD4^{+}CD25^{+}GITR^{+}V\alpha2^{+}V\beta5.1^{+}$ Tregs sorted from OTII mice, in the presence of $OVA_{323\text{-}339}$. Cells were stained for actin, TCR and PD-1, interactions are observed on filopodia extending from $\alpha$-$SMA^{+}$ CAFs. Related to Figure 3. 30fps, scale bar: 1μm.

**Pipeline for analysis of time-lapse videos**

1. Identify Treg tracks using “Trackmate” plugin. Call the output “Tregs”.

2. Find max value, M, in “Tregs” stack:

Image → Stack → z-Project, then

Analyse → Measure → Min & max grey value

3. Identify CAF cells in raw time-lapse image (requires “PHANTAST” plugin). Call the output “CAFs”:

a. File → Save as → Image Sequence

b. Process → Batch → Macro. Run the following commands

run("32-bit"); run("PHANTAST", "sigma=10 epsilon=0.06 new"); run("Analyse Particles...", "size=5000-Infinity show=Masks"); run("Invert LUT");

c. File -> Import -> Image Sequence (to convert individual images to stack)

d. Manual corrections: Delete region, Separate cells (draw background-coloured, 3-pixel wide regions between cells)

e. Identify 3d CAFs from segmented image:

Analyse → 3D-object Counter

f. Image → Type → 32bit

4. Multiply “CAFs” by 10^N, where N such that 〖10〗^N > M: Process → Math → Multiply → 10^N

(32bit images allow pixel values up to 2^32).

5. Create an image with pixel values equal to the minimum value between the corresponding pixels in the “CAFs” and “Tregs” images. Call the output “Intersected”. Because of (2, 4), the value at intersecting pixels is that of the respective T-reg. cell. If a Treg cell never intersects any CAF cell

the minimum between the value of any of its pixels and the corresponding pixels of the CAF image will be zero throughout the time lapse: Process → Image calculator → "Image 1": Tregs, "Operation": Min, "Image 2": CAFs.

6. Add the "CAFs" and "Intersected" images. Call the output "Added". Remark: In the "Added" image a pixel value of 3000472 indicates that the T-reg with value (label) 472 in the "T-regs" time-lapse intersects the CAF cell with label 3000000 (in this case we have multiplied with 10^6 in step 4). Therefore, the number of times the value 3000472 appears throughout the time lapse is the total number of pixels that this CAF-Treg intersection occupies throughout the time lapse: Process → Image calculator → "Image 1": CAFs, "Operation": Add, "Image 2": Intersected.

7. In this and the next two steps we reduce the CAF-T-reg. intersection areas to single pixels per time point. Then, the number of times the value, say, 3000472 appears throughout the time lapse will reveal the number of time points T-reg 472 intersects the CAF cell 3000000, which is the statistic of interest. First, use the "Find Maxima" command on the "Added" image, which returns a single pixel for each local maximum, that is, for each region of intersection (because of step 6). The result is a binary image with value 255 on pixels corresponding to such maxima and 0 elsewhere:

a. File → Save as → Image Sequence

b. Process → Batch → Macro. Run the following commands: run("Find Maxima...", "prominence=10 output=[Single Points]"); run("32-bit");

c. File → Import → Image Sequence (to convert individual images to stack)

8. Multiply the image from the previous step with any value ≥10^(N+1), where N, the value used in step (4). Call the output "Maxima": Process → Math → Multiply → 10^(N+1).

9. Find “minimum” image (as in step 5) between “Added” and “Maxima”. Because of (6, 7, 8), the output is an image with isolated non-zero pixels with values corresponding to the CAF-Treg intersection values from the “Added” image. Call the output “#_Histogram”, where # is some number: Process → Image calculator → “Image 1”: Added, “Operation”: Min, “Image 2”: Maxima.

10. Create histogram for “#_Histogram” image. Based on the above steps, the height of the bar at the value x (e.g., 3000472) indicates the number of time-points the CAF-Treg intersection with value x appears throughout the time-lapse. Find maximum value, “MAX”, in “#_Histogram” image: Image → Stack → z-Project, then Analyse → Measure → Min & max gray value. Analyse → Histogram: set "Bins" and "Xmax" equal to “MAX”, deselect "Use pixel value range", select "Stack histogram".

**Mouse Genotyping**

Mice were genotyped according to previously published protocols regarding the αSMA-tk mice and αSMAcre mice from Jackson Laboratories (Bar Harbor, ME, USA) and the Atg5fl/fl mice from RIKEN BioResource Center (Koyadai,Tsukuba-shi, Ibaraki, Japan). The following primers were used (Invitrogen):

For Atg5fl/fl mice:

- exon 3-1: 5’- GAA TAT GAA GGC ACA CCC CTG AAA TG -3’
- short2: 5’- GTA CTG CAT AAT GGT TTA ACT CTT GC -3’
- check2: 5’- ACA ACG TCG AGC ACA GCT GCG CAA GG -3’

For αSMAcre mice:

- transgene R: 5'- ACA TGT CCA TCA GGT TCT TGC -3'
- internal positive control F: 5'- AGT GGC CTC TTC CAG AAA TG -3'
- internal positive control R: 5'- TGC GAC TGT GTC TGA TTT CC -3'
- transgene F: 5'- GGT GTT AGT TGA GAA CTG TGG AG -3'

For αSMA-tk mice:

- transgene R: 5'-GTG GTG GTT TTC CCC ATC C -3'
- internal positive control F: 5'- AGT GGC CTC TTC CAG AAA TG -3'
- internal positive control R: 5'- TGC GAC TGT GTC TGA TTT CC -3'
- transgene F: 5'- ACC AAG AAC CCT GTC TGT GG -3'

**TEM additional information**

Receipt of resin embedding media

-Durcupan single component A, M epoxy resin 10 ml

-Durcupan single component B, hardener 964 10ml

-Durcupan single component C, accelerator 960 0,3ml

-Durcupan single component D, plasticiser 0,3ml
